# Nerve Excitability Differences in Slow and Fast Motor Axons of the Rat: more than just *I*_*h*_

**DOI:** 10.1101/613984

**Authors:** James M. Bell, Chad Lorenz, Kelvin E. Jones

## Abstract

**Objective:** The objective was to determine if choice of anaesthetic confounded previous conclusions about the differences in nerve excitability indices between fast and slow motor axons.

**Methodology:** Nerve excitability of the rat sciatic nerve was tested while measuring responses of motor axons innervating the slow-twitch soleus (SOL) and fast-twitch tibialis anterior (TA) muscles. The experiments were conducted with sodium pentobarbital (SP) anaesthetic and compared to previous results that used ketamine-xylazine (KX).

**Results and Conclusions:** Previous conclusions about the differences in nerve excitability indices between TA and SOL motor axons using KX were corroborated and extended when experiments were done with SP. Nerve excitability indices sensitive to changes in hyperpolarization-activated inwardly rectifying cation current (*I*_*h*_) indicated an increase in *I*_*h*_ in SOL axons compared to TA axons (*e.g*. S3 (−100 %), t=7.949 (df=10), *p* < 0.0001; TEh (90–100 ms), t=2.659 (df=20), *p* = 0.0145; hyperpolarizing I/V slope, t=4.308 (df=19), *p* = 0.0004). SOL axons also had a longer strength-duration time constant (t=3.35 (df=20), *p* = 0.0032) and a longer and larger magnitude relative refractory period (RRP (ms) t=3.53 (df=12), *p* = 0.0041; Refractoriness at 2 ms t=0.0055 (df=9), *p* = 0.0055).

Anaesthetic choice affected many measures of peripheral nerve excitability with differences most apparent in tests of threshold electrotonus and recovery cycle. For example, recovery cycle with KX lacked a clear superexcitable and late subexcitable period. We conclude that KX had a confounding effect on nerve excitability results consistent with ischaemic depolarization. Results using SP revealed the full extent of differences in nerve excitability measures between putative slow and fast motor axons of the rat. These differences have important implications for the use of nerve excitability measures during processes such as ageing where it is believed that there is a selective loss of fast axons.

**New & Noteworthy:** Nerve excitability testing is a tool used to provide insight into the properties of ion channels in peripheral nerves. It is used clinically to assess pathophysiology of motor axons. Researchers customarily think of motor axons as homogeneous; however, we demonstrate there are clear differences between *fast* and *slow* axons in the rat. This is important for interpreting results with selective motor neuronopathy, like aging where *fast* axons are at high risk of degeneration.

## 1. Introduction

One of the long-standing axioms of mammalian motor neuroscience is that there is a functionally relevant association between the intrinsic properties of motor neurons and the contractile physiology of the muscle units they innervate [9, 12, 38]. For example, in the healthy adult rat the faster the time-course of the afterhyperpolarization (AHP), the faster the axonal conduction velocity and the faster the time-course of the twitch tension [14, 15]. However, conduction velocity alone is a poor measure of the intrinsic properties of motor axons and the motor neurons they arise from, as it is primarily correlated with anatomical measures [11, 36]. In a previous study we used nerve excitability testing to infer differences between the intrinsic properties of motor axons innervating physiologically distinct target muscles. For convenience we labeled motor axons to soleus (SOL) as slow and those innervating tibialis anterior (TA) as fast since rat SOL muscle is composed of about 80 % slow-twitch [17] while about 94 % of TA is histologically fast-twitch [10]. The primary finding was that fast motor axons have a reduced hyperpolarization-activated inwardly rectifying cation current (*I*_*h*_) [29]. In that study the anaesthetic was ketamine-xylazine (KX). However, pilot studies evaluating sodium pentobarbital (SP) as a potential substitute produced very different results, so use of SP was abandoned and the experiments were completed using KX.

Many previous studies used KX for nerve excitability testing in rodents (see Supplementary Table 1 for details and references). Of the 38 studies, 19 tested the hindlimb, but only 2 of those used KX. One was our previous study, which had results with an uncharacteristic shape most noticeable in the recovery cycle and threshold electrotonus [29]. Another recent study had similarly uncharacteristic results [33]. In the tail, KX has been used with no apparent anomalies [13, 16, 31].

In this study, we gathered additional data using SP as the primary anaesthetic to determine if the anaesthetic effect was reproducible and to determine if differences between fast and slow motor axons remained under a different anaesthetic. We hypothesized (1) that the switch in anesthetics would not alter the excitability of peripheral nerves, given that the primary effects of the anaesthetics are at the synapse, and (2) that the differences between fast and slow axons would be similar to previous reports.

## 2. Material and Methods

All animal studies were conducted in accordance with the Canadian Council on Animal Care Guidelines and Policies with approval from the Animal Care and Use Committee: Health Sciences for the University of Alberta. Approval was granted for the use of 6 additional animals for these experiments. We duplicated the methods used in our previous study [29], with one important exception: sodium pentobarbital replaced ketamine-xylazine as the anaesthetic. The dosage was 40 mg/kg sodium pentobarbital, with additional doses as needed during the experiment. The results from both legs of female Sprague Dawley rats, mass 279 ± 12 g, 13 weeks old, are presented. The following section is a summary of the materials and methods which were previously described in detail.

### Surgery and Recordings

Nonpolarizing stimulating electrodes were used for percutaneous stimulation of the sciatic nerve following local hair removal. We denervated the extensor digitorum longus, flexor digitorum longus, medial gastrocnemius, and plantaris to ensure that the recorded compound muscle action potential (CMAP) in the target muscle was not confounded by a CMAP in any other muscles. Since soleus and lateral gastrocnemius axons share a nerve branch, the lateral gastrocnemius could not be denervated, so the distal 60 % was excised. We then inserted stainless steel wire hooks into the muscles of the leg for electromyographic (EMG) recordings. Measurements were first taken from the TA and then from the SOL. The rats were placed on a warming pad and core temperature monitored and controlled. Incisions for placement of EMG recordings were closed with staples to reduce heat loss from the lower limbs. For each rat, the time from anaesthetic administration to the completion of surgery and all recordings was less than one hour. The rats were then euthanized by anaesthetic overdose and cervical dislocation.

To measure the excitability of TA and SOL axons, QTRAC software (protocol “TRONDCMW”) was used to control stimulation in a closed-loop feedback control system with EMG recordings. This protocol, described in more detail by Kiernan and Lin [27], provides five measures of nerve excitability: recovery cycle, threshold electrotonus, rheobase, strength-duration time constant, and current-threshold. Recovery cycle provides a measure of the nerve’s excitability after brief supramaximal conditioning stimuli at varying delays. Threshold electrotonus measures the nerve’s response to long-lasting (100 ms) sub-threshold depolarizing and hyperpolarizing currents at varying delays. Rheobase and the strength-duration time constant describe the relationship between the intensity and duration of a threshold stimulus. The current-threshold relationship measures threshold changes after a long (200 ms) subthreshold current pulse over a range of polarities. In order to specifically look at the role of *I*_*h*_, additional 200 ms and 300 ms hyperpolarizing threshold electrotonus measures were also included [32].

### Analysis

The data were analyzed using parametric (t-test) or non-parametric (Mann-Whitney U-test) tests as appropriate. Data were not collected for every measure from every animal, ruling out the use of paired statistical tests, so independent tests were used and degrees of freedom are given in Table 1. Missing data for nerve excitability tests is an important topic that we are addressing for large human data sets [4]. We recognize that, given the missing data issue which precluded using paired tests, there is an increased risk of type 2 errors in the analysis. However, the primary findings of differences in *I*_*h*_ between fast and slow axons are present and independent of the type of statistical test used to generate the p-value. We have elected to report raw p-values rather than use an arbitrary threshold for indicating significance [2]. Summary statistics are presented as mean or median (standard deviation, SD, or interquartile range, IQR). Graphs display the mean and standard error of the mean unless stated otherwise.

**Table 1:**
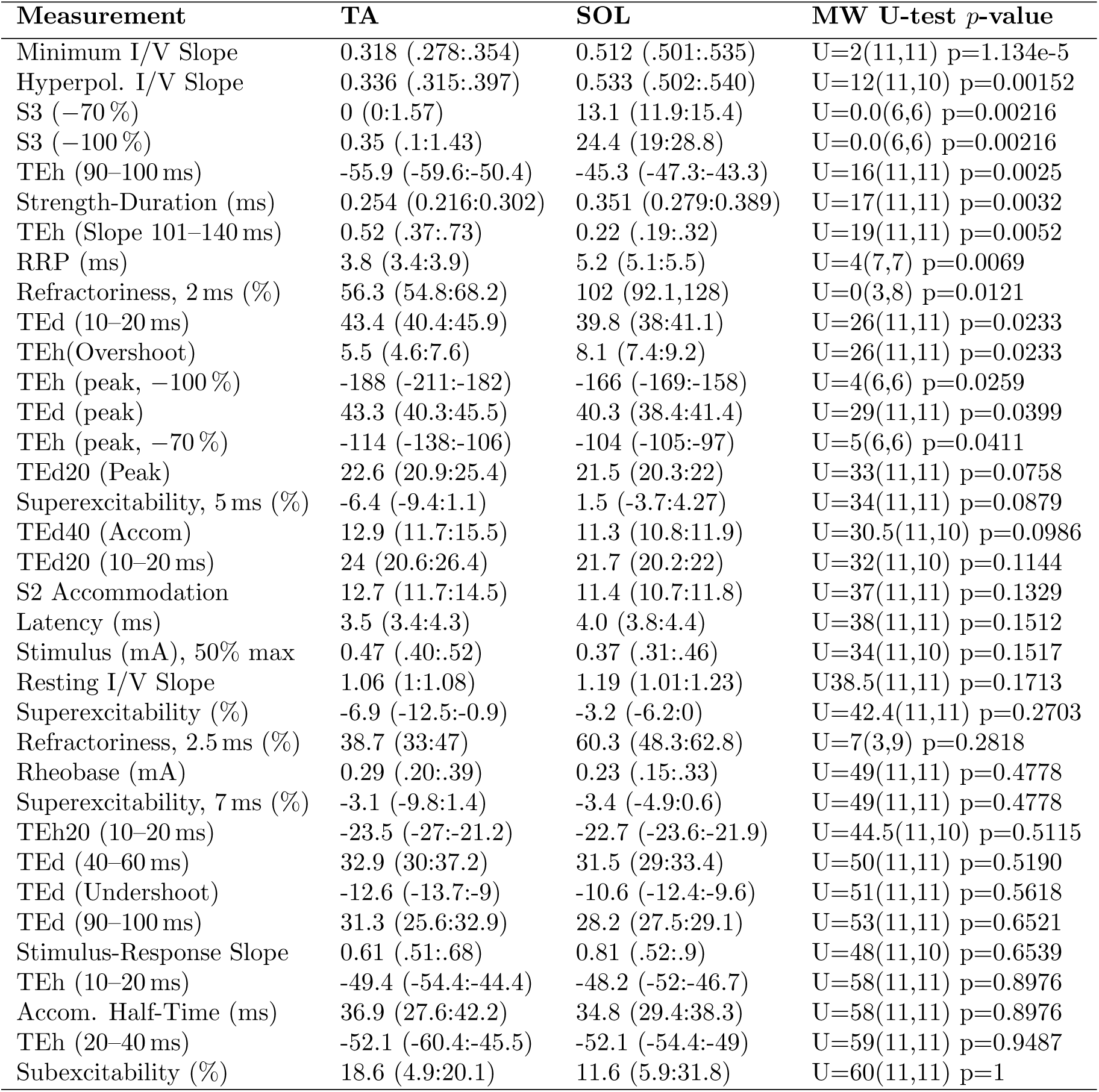
Median (and IQR) for standard measures of nerve excitability, comparing the response of fast (tibialis anterior, TA) and slow (soleus, SOL) motor axons. Mann-Whitney U-test statistic (with numbers in each group) and associated p-value were used to rank order the comparisons.

## 3. Results

There was a clear and consistent effect of anesthetic on measures of nerve excitability, contrary to our first hypothesis. Figure 1 shows comparison of the anaesthetic effects for both TA and SOL axons, with some differences in nerve excitability indices highlighted in graph insets. The strength-duration results for both axons showed a decrease in rheobase for KX (Supplementary Figures 1a and 1b). The recovery cycle results were the most clearly affected, with absent superexcitable and late subexcitable periods with KX. During threshold electrotonus using KX, the S1 phase was absent (measured by the 10–20 ms and peak TEd values) and the threshold change during hyperpolarizing currents was attenuated. A supplementary table 2 includes quantitative comparisons of these key differences.

**Figure 1:**
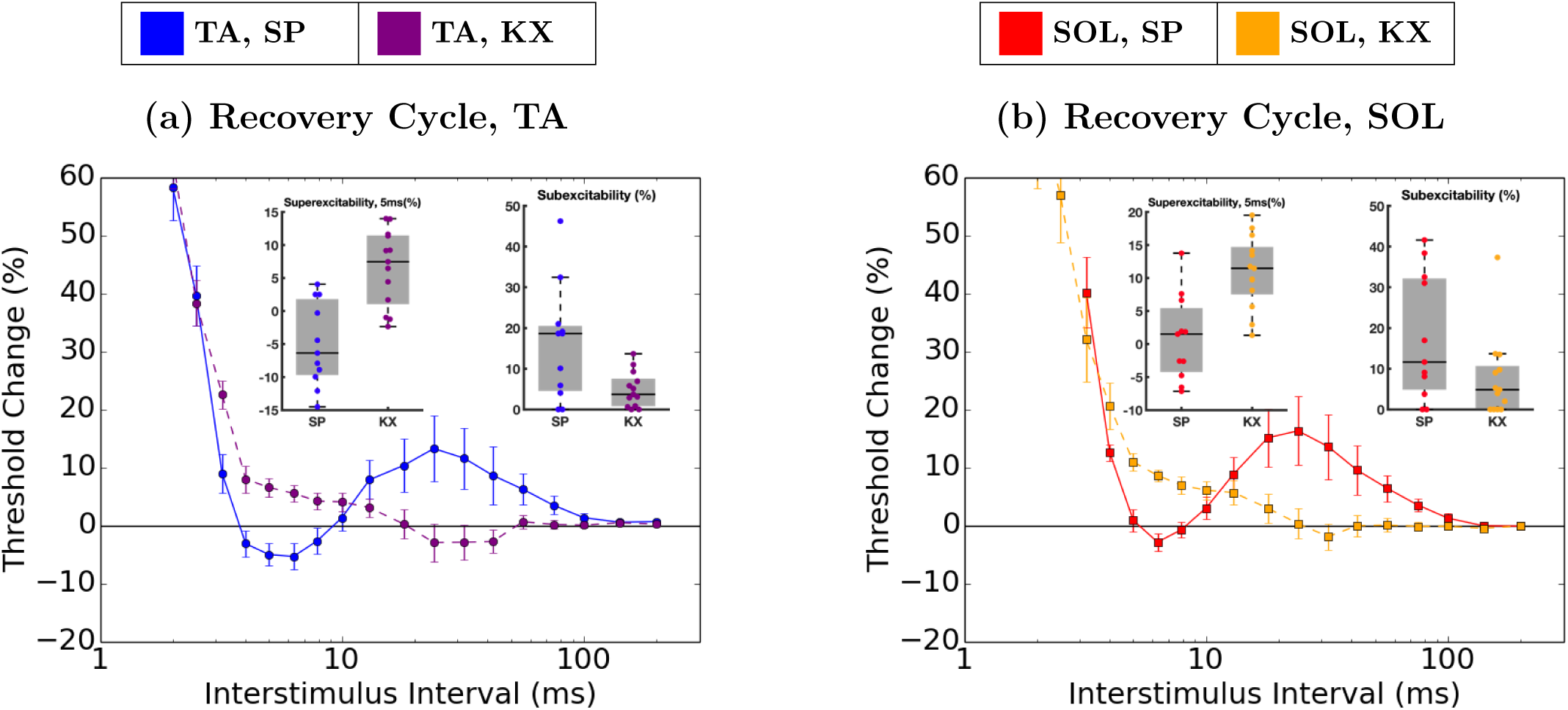
Anaesthetic effects of ketamine-xylazine (KX) compared to sodium pentobarbital (SP) on the motor axons of tibialis anterior (TA) and soleus (SOL). Mean values are plotted, with standard error bars. For Strength Duration and Threshold Electrotonus figures, see the supplementary material. The inset graphs illustrate the median (solid red line), IQR (gray box) and 1.5*IQR (whiskers).

The differences between TA and SOL axons using SP as the anaesthetic were similar to but more extensive than those previously reported using KX [29], as shown in Figure 2. Differences in some key nerve excitability indices are illustrated in the graph inserts. In recovery cycle, the TA axons had a shorter relative refractory period (RRP), were less refractory at short latency and had a larger early superexcitable period (though at the 5 ms time point, nonparametric significance testing was equivocal, Figure 2b). For depolarizing threshold electrotonus, the S1 phase was absent for SOL (TEd at 10–20 ms, Figure 2c); hyperpolarizing threshold electrotonus was reduced for SOL (TEh at 90–100 ms, Figure 2c). Threshold I/V tests (Figure 2e) showed that the hyperpolarizing and minimum I/V slopes were larger for SOL. The strength-duration time constant was shorter for TA (*p* <0.001). The stimulus-response curve was not notably different (Figure 2f). Table 1 includes quantitative comparisons of these differences.

**Figure 2:**
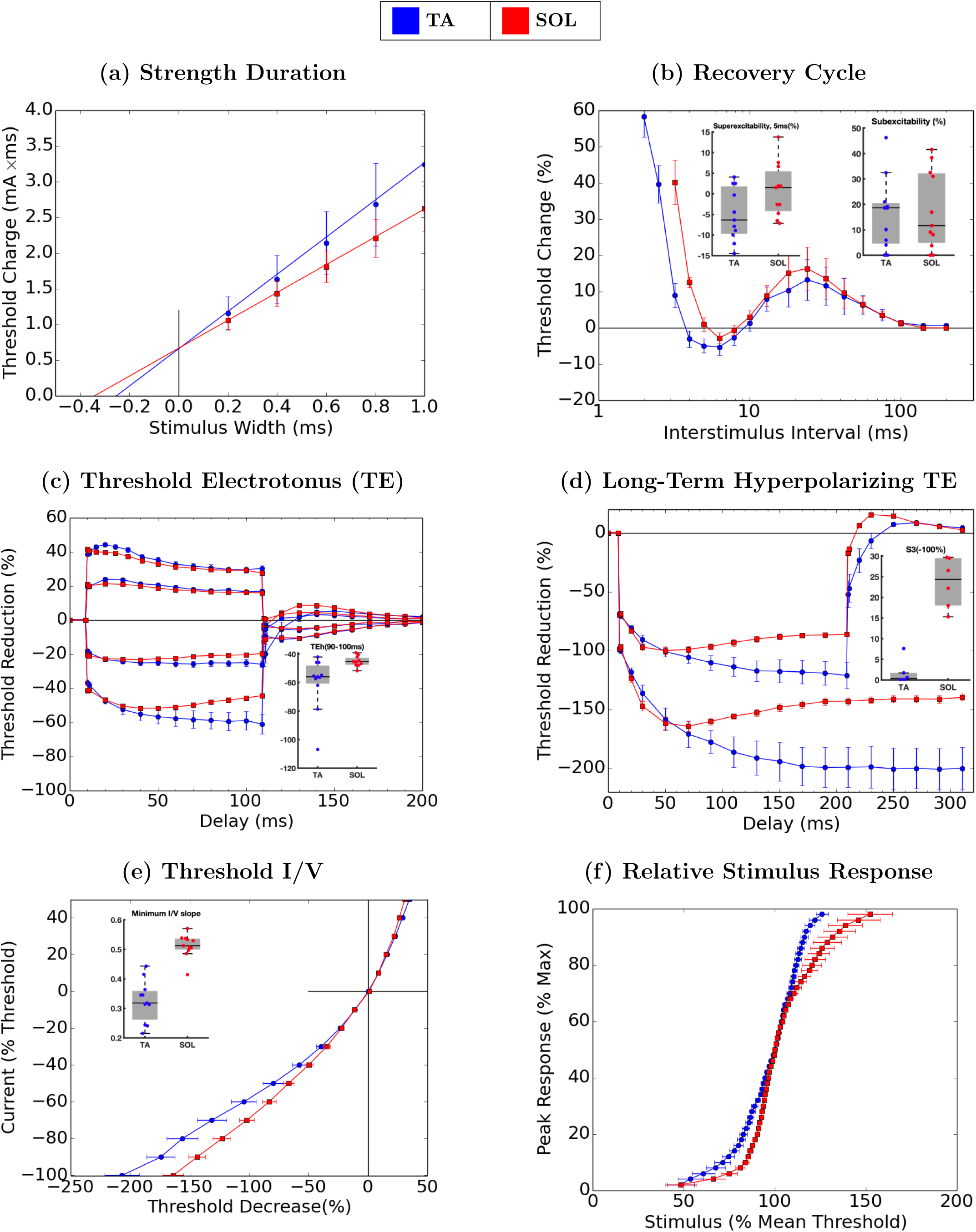
Differences between fast (tibialis anterior, TA) and slow (soleus, SOL) motor axons. Mean values are plotted, standard error bars.

New to this study, we present a comparison of longterm hyperpolarizing threshold electrotonus (Figure 2d). It shows a significant difference in the S3 phase, which measures the ratio of peak threshold increase to the value in the 10 ms before the end of the hyperpolarizing conditioning current [32]. The error bars in the plot illustrate that error in the estimate of the mean was greater for TA compared to SOL axons.

## 4. Discussion

### 4.1. Confounding Effects of Ketamine-Xylazine

The results of Figure 1 show a clear anaesthetic effect. Our SP results are consistent with the literature: electrodiagnostic nerve test results in most nerves have a characteristic shape, regardless of the anaesthetic used. The one exception is when KX is used when testing the sciatic nerve (e.g. [29, 33]). We suggest that the most parsimonious explanation of these results is that blood flow to the nerve is reduced due to the vasoconstrictive effect of the alpha-adrenergic agonist xylazine [25], causing ischaemia which results in depolarization. According to Kiernan and Bostock [26], ischaemia results in decreased rheobase (Supplementary Figures 1a and 1b); absent or diminished superexcitable and late subexcitable periods (Figures 1a and 1b); a characteristic fanning-in during threshold electrotonus (Supplementary Figures 1e and 1f); and an increase in resting I/V slope (not shown in a figure, but see Supplementary Table 2). The results using KX are consistent with expected changes arising from ischaemia; however, we did not observe the expected increase in the strength-duration time constant (Supplementary Figures 1a and 1b) or minimum threshold I/V slope (not shown), which Kiernan and Bostock [26] reported to be altered during ischaemia. KX appears to have a ischaemia-like effect, though some unexplained differences remain.

Ketamine is less likely than xylazine to give rise to the observed changes in nerve excitability indices. Ketamine’s anaesthetic effect is attributed to NMDA receptors, which are negligible on the peripheral nerves [22]. At high doses, local injection of ketamine blocks nerve Na^+^ channels. However, the anaesthetic effects reported here with KX are not consistent with reduced Na^+^ influx, and the circulating concentration of ketamine following i.p. injection is below that required for a blocking effect on Na^+^ channels [22, 39].

Xylazine has been used in many previous studies of nerve excitability in rodents, apparently without the same ischaemia-like effects demonstrated in the motor axons in the sciatic nerve. We suggest that the off-target vasoconstrictive ischaemic effects on the sciatic nerve are more pronounced than on other nerves because the sciatic nerve has a poorer nutrient artery supply compared to other nerves. While there is little literature supporting that assertion, it is commonly believed among anaesthesiologists who do peripheral nerve blocks for regional anaesthesia. For example, Warman et al. [35] says that many anaesthesiologists avoid using vasoconstrictors when injecting around the sciatic nerve “due to a perceived poor blood supply” [35, p. 32]. We suggest that further research in this area would be of interest to researchers and practitioners alike, but the purpose of this paper is to provide updated results for differences between fast and slow motor axons.

### 4.2. Differences Between Slow and Fast Motor Axons

In spite of the confounding effects of the anaesthetic used in our previous study [29], the conclusions are compatible with the present study; a primary difference between slow SOL and fast TA motor axons is likely due to reduced *I*_*h*_ in the fast motor axons. We provide Figure 2 as an improved representation of those differences. A number of the nerve excitability measures reported in Table 1 are sensitive to changes in *I*_*h*_, including the top 5 measures ranked in order of p-values. Axons with greater levels of *I*_*h*_ conductance are expected to show greater minimum and hyperplorizing I/V slopes, greater values for S3, and smaller threshold reduction at the end of the 100 ms hyperpolarizing threshold electrotonus. Lorenz and Jones [29] observed that the threshold reduction at 90–100 ms during a hyperpolarizing threshold electrotonus test was greater in TA presumably because of reduced *I*_*h*_, a conclusion supported by previous tests with cesium, an HCN-blocker, which caused similar effects [3, 5, 37]. In the present results, that difference was even more pronounced. The additional 200–300 ms hyperpolarizing threshold electrotonus and S3 measures further supports this conclusion, since it is strongly dependent on *I*_*h*_ [32].

Lorenz and Jones [29] also observed significant but small-magnitude differences in rheobase and hyperpolarizing threshold electrotonus at 20–30 ms, changes which are not explained by differences in *I*_*h*_ and not reproduced in the current data. Our updated results suggest that in addition to *I*_*h*_, there are differences in the intrinsic properties of fast and slow axons associated with the strength-duration time constant and the early refractory period measured during the recovery cycle (Figure 2b). The strength-duration time constant is associated with the amount of low threshold persistent Na^+^ conductance and passive electrotonic properties [6]; thus our results suggest an increase in *I*_*N*_*a*_*p*_ and/or a longer membrane time constant in slow motor axons. The increased duration of the relative refractory period (RRP) in slow axons could arise in part from the increase in *I*_*N*_ *a*_*p*_ which tends to slow the recovery of transient Na^+^ conductance from inactivation. However, there are a number of other intrinsic properties that contribute to the shift from the RRP to the superexcitable period that could be implicated, including the current arising from the capacitance of the internode which generates the depolarizing afterpotential [27]. Our results are not sufficient to give a mechanistic explanation for the differences in the early recovery cycle measures between fast and slow axons. It is clear that our results can not be explained by changes in *I*_*h*_ alone.

### 4.3. Functional Implications of Differences in *I*_*h*_

The measurement of the S3-phase following long duration hyperpolarization is reminiscent of the sag process originally described in the soma of cat spinal motor neurons by Ito and Oshima [24]. It was later determined that the duration of the afterhyperpolarization (AHP) in cat motor neuron somata was associated with the amount of sag (now known to arise from *I*_*h*_); shorter duration AHPs in fast motor neurons are partly a consequence increased *I*_*h*_ that counteracts Ca-dependent K^+^ conductance underlying the AHP [19]. This makes functional sense as fast motor neurons which are associated with faster twitch muscle units require a higher rate of discharge to generate fused contractions, and a faster time-course of the AHP enables higher discharge rates [14, 15]. However, the present results suggest that *I*_*h*_ in fast is *decreased* relative to slow motor axons. This implies that regulation of expression of HCN channels, which generate *I*_*h*_, would be different in the soma versus axon, a suggestion that is in keeping with current evidence [1].

The functional role of *I*_*h*_ in the cell body of neurons has been discussed in terms of regulating rhythmic discharge properties (i.e. the AHP) and facilitating pacemaker-like activity (e.g. [30]). Not surprisingly, dysfunction in *I*_*h*_ is linked to disorders of rhythmic discharge like seizures [7]. It is clear that *I*_*h*_ is present in myelinated and unmyelinated axons of a wide range of species [8, 28], but what is the functional role of *I*_*h*_ in myelinated motor axons? One proposition is that *I*_*h*_ may limit activity-dependent hyperpolarization of the axons and the accompanying branch point failure [3, 23, 32, 34]. Rat TA and SOL motor axons experience very different patterns of impulse traffic in functional activities. Soleus motor axons must faithfully conduct 300-500 thousand impulses per day in bursts of activity that last up to seven minutes for 5-8 hours a day, though the average discharge rate during a burst is modest [20]. In contrast TA motor axons conduct brief, high-frequency bursts of about 4-5 impulses per step followed by long periods of rest [18]. Using published data from the fast extensor digitorum longus muscle of the rat, in place of tibialis anterior, these axons must conduct 3-11 thousand impulses per day in short, high-frequency bursts [20]. It seems plausible that the increased *I*_*h*_ in slow SOL motor axons of the rat may be an activity-dependent adaptation to the pattern of impulses normally carried by these axons that would circumvent activity-dependent hyperpolarization.

## 5. Conclusions

The use of ketamine-xylazine, a common anaesthetic for nerve excitability tests, was shown to have a confounding effect in the rat sciatic nerve. Nerve excitability test results using sodium pentobarbital are superior and show the full extent of differences between fast and slow motor axons.

Previous results suggesting increased *I*_*h*_ in slow motor axons were confirmed more clearly in the absence of the confounding effect of anaesthetic. We purposefully chose divergent populations of motor axons to assess whether nerve excitability tests results in healthy young adult rats were different. When interpreting the results of a nerve excitability test it will be important to consider the proportion of slow:fast axons in the nerve being tested. In aged rats, when there is increased degeneration of fast axons [21], one should expect the remaining axons to appear more like slow axons. Similarly, if expression of *I*_*h*_ in motor axons adapts to activity-dependent changes we would expect decreases in axonal impulse traffic to promote adapation towards the fast axon phenotype described here.

## Acknowledgments

The authors are grateful to Neil Tyreman for his expert technical assistance in the lab. Funding: This work was supported by grants (KJ): NSERC Discovery Grant [RGPIN-2017-05624], CIHR Neuromuscular Research Partnership with Muscular Dystrophy Canada, and the ALS Society of Canada [201003JNM-225975-MOV]; and scholarships (JB): the University of Alberta Faculty of Medicine and Dentistry Graduate Recruitment Scholarship, a Queen Elizabeth II Graduate Student Scholarship and a NSERC-Alexander Graham Bell Canada Graduate Scholarship.

