## Supplemental Table and Figures for "Nerve Excitability Differences in Slow and Fast Motor Axons of the Rat: more than just *I*_*h*_"

### Supplementary Figures and Analysis for: Nerve Excitability Differences in Slow and Fast Motor Axons of the Rat: more than just $I_h$

James M. Bell[1,2], Chad Lorenz[1,3] and Kelvin E. JonesE*[1,3]

[1]Neuroscience and Mental Health Institute, University of Alberta
[2]Department of Computing Science, University of Alberta
[3]Faculty of Kinesiology, Sport, and Recreation, University of Alberta

April 24, 2019

| **Species** | **Location** | **Anaesthetic** | **Reference** |
|---|---|---|---|
| Rat | Sciatic | KX | [17, 36] |
| Rat | Forelimb | Hypnorm/Midazolam | [3] |
| Rat | Tail | Isoflurane | [24] |
| Rat | Tail[1] | KX | [14, 15, 33] |
| Mouse | Sciatic | Fentanyl/Droperidol/Midazolam | [18] |
| Mouse | Sciatic | Hypnorm/Midazolam | [1, 2, 20–23, 30–32] |
| Mouse | Sciatic | Isoflurane | [4, 6, 11, 12] |
| Mouse | Sciatic | Pentobarbital | [35] |
| Mouse | Tail | Isoflurane | [5, 8, 10, 13, 16, 19, 25–29, 34] |
| Mouse | Tail | Medetomidine/Midazolam/Butorphanol | [29] |
| Mouse | Tail | pentobarbital | [38] |
| Mouse | Tail/Sciatic | Isoflurane | [7, 9, 37] |

Table 1: A list of anaesthetics used for previous electrodiagnostic nerve testing in rodents. Note 1: One study [14] included a variety of tests focused on the sciatic nerve; the electrodiagnostic nerve test stands out as the only test in that study which was performed on the tail instead.

*

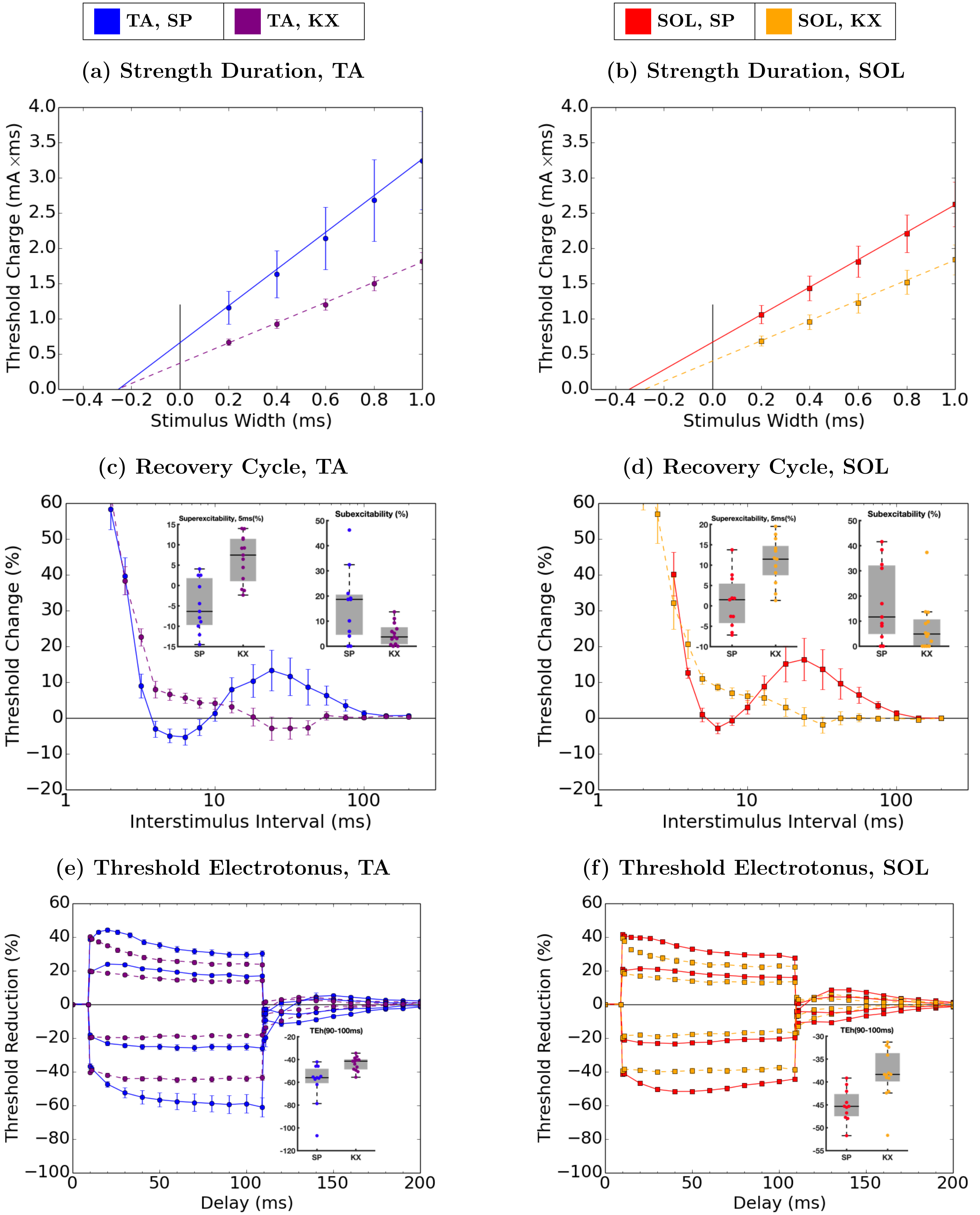


Figure 1: Anaesthetic effects of ketamine-xylazine (KX) compared to sodium pentobarbital (SP) on the motor axons of tibialis anterior (TA) and soleus (SOL). Mean values are plotted, with standard error bars.

| **Measurement** | **SP, SOL** | **KX, SOL** | ***p*, SOL** | **SP, TA** | **KX, TA** | ***p*, TA** |
|---|---|---|---|---|---|---|
| Strength-Duration (ms) | 0.34 (0.02) | 0.28 (0.02) | 0.03 | 0.25 (0.02) | 0.25 (0.02) | 0.89 |
| Rheobase (mA) | 1.8 (1.1) | 1.3 (1.1) | 0.05 | 2.2 (1.2) | 1.4 (1.1) | 0.01 |
| RRP (ms) | 5.2 (1.1) | 12.6 (1.5) | 0.02 | 3.8 (1.1) | 6.0 (1.3) | 0.03 |
| Refractoriness, 2 ms (%) | 113 (21) | 99 (16) | 0.63 | 58 (6) | 64 (7) | 0.61 |
| Refractoriness, 2.5 ms (%) | 54 (9) | 57 (8) | 0.82 | 40 (5) | 37 (5) | 0.73 |
| Superexcitability, 5 ms (%) | 0.9 (2.0) | 10.9 (1.5) | <0.001 | -5.0 (1.9) | 6.5 (1.6) | <0.001 |
| Superexcitability, 7 ms (%) | -1.8 (1.3) | 7.8 (1.1) | <0.001 | -4.0 (2.2) | 5.2 (1.3) | 0.001 |
| Superexcitability (%) | -1.9 (4.5) | 4.7 (5.2) | 0.003 | -5.4 (6.7) | 2.5 (5.1) | 0.004 |
| Subexcitability (%) | 17.6 (4.7) | 7.6 (2.8) | 0.07 | 16.0 (4.3) | 4.8 (1.2) | 0.01 |
| TEd (10–20 ms) | 39.0 (1.1) | 29.4 (1.6) | <0.001 | 43.0 (1.2) | 32.8 (1.2) | <0.001 |
| TEd (peak) | 39.8 (1.0) | 31.6 (1.3) | <0.001 | 43.1 (1.0) | 35.0 (1.0) | <0.001 |
| TEd (40–60 ms) | 31.3 (1.4) | 23.8 (1.5) | 0.002 | 33.0 (1.8) | 25.6 (1.2) | 0.002 |
| Accom. Half-Time (ms) | 35.9 (2.9) | 26.8 (4.0) | 0.09 | 36.4 (3.2) | 26.1 (2.0) | 0.009 |
| TEd (90–100 ms) | 28.2 (0.9) | 22.6 (1.0) | <0.001 | 29.8 (1.7) | 23.8 (1.0) | 0.005 |
| TEd40 (Accom) | 11.33 (0.44) | 10.07 (0.65) | 0.13 | 14.14 (1.46) | 11.41 (0.55) | 0.07 |
| TEd20 (10–20 ms) | 21.0 (0.5) | 16.2 (0.9) | <0.001 | 23.7 (1.1) | 18.0 (0.7) | <0.001 |
| TEd20 (Peak) | 21.1 (0.4) | 17.0 (0.7) | <0.001 | 23.4 (0.9) | 18.6 (0.6) | <0.001 |
| S2 Accommodation | 11.53 (0.46) | 9.04 (0.71) | 0.01 | 13.29 (1.29) | 11.17 (0.55) | 0.12 |
| TEd (Undershoot) | -10.9 (0.5) | -9.1 (0.5) | 0.02 | -11.6 (1.3) | -10.2 (0.6) | 0.31 |
| TEh (10–20 ms) | -48.8 (1.0) | -39.6 (1.1) | <0.001 | -50.0 (2.2) | -43.1 (1.2) | 0.01 |
| TEh (20–40 ms) | -51.5 (1.2) | -39.9 (1.4) | <0.001 | -55.0 (3.8) | -44.1 (1.4) | 0.008 |
| TEh (90–100 ms) | -45.2 (1.1) | -38.3 (1.5) | 0.002 | -60.1 (5.5) | -43.8 (1.7) | 0.006 |
| TEh (Slope 101–140 ms) | 0.246 (0.034) | 0.052 (0.050) | 0.005 | 0.575 (0.114) | 0.128 (0.033) | <0.001 |
| TEh20 (10–20 ms) | -22.94 (0.65) | -18.69 (0.45) | <0.001 | -24.02 (1.21) | -19.74 (0.48) | 0.002 |
| TEh(Overshoot) | 8.36 (0.46) | 6.01 (0.66) | 0.01 | 6.04 (0.83) | 4.28 (0.47) | 0.06 |
| Hyperpol. I/V Slope | 0.524 (0.016) | 0.467 (0.026) | 0.08 | 0.358 (0.036) | 0.407 (0.026) | 0.28 |
| Resting I/V Slope | 1.177 (0.056) | 1.468 (0.115) | 0.04 | 1.037 (0.044) | 1.275 (0.062) | 0.006 |
| Minimum I/V Slope | 0.512 (0.012) | 0.456 (0.024) | 0.07 | 0.323 (0.021) | 0.391 (0.025) | 0.06 |
| Peak Response (mV) | 4.3 (1.5) | 3.0 (1.3) | 0.47 | 4.8 (1.8) | 1.7 (1.1) | 0.06 |
| Stimulus (mA), 50% max | 2.5 (1.1) | 1.8 (1.1) | 0.05 | 3.2 (1.2) | 1.9 (1.1) | 0.002 |
| Stimulus-Response Slope | 4.9 (1.2) | 4.5 (1.2) | 0.67 | 4.6 (1.2) | 4.1 (1.1) | 0.55 |
| Latency (ms) | 4.10 (0.14) | 4.35 (0.20) | 0.35 | 3.85 (0.21) | 4.29 (0.30) | 0.26 |

Table 2: Mean (and standard deviation) for some key differences in nerve excitability, comparing the effects of anaesthetics ketamine-xylazine (KX) and sodium pentobarbital (SP) in tibialis anterior (TA) and soleus (SOL) motor axons. The columns of raw p-values were generated using t-tests and are reported without correction for multiple comparisons. The calculation for *Superexcitability (%)* did not use the standard QTRAC output because the QTRAC output is not correct when the superexcitable period is not present. Instead, the value was calculated as the minimum 3-sample moving average in the first 8 samples (i.e. up to 12ms).
